# *In vivo* stabilization of endogenous chloroplast RNAs by customized artificial pentatricopeptide repeat proteins

**DOI:** 10.1101/2020.11.21.392746

**Authors:** Nikolay Manavski, Louis-Valentin Meteignier, Margarita Rojas, Andreas Brachmann, Alice Barkan, Kamel Hammani

## Abstract

Pentatricopeptide repeat (PPR) proteins are helical repeat-proteins that bind RNA in a modular fashion with a sequence-specificity that can be manipulated by the use of an amino acid code. As such, PPR repeats are promising scaffolds for the design of RNA binding proteins for synthetic biology applications. However, the *in vivo* functional capabilities of artificial PPR proteins built from consensus PPR motifs are just starting to be explored. Here, we report *in vivo* functions of an artificial PPR protein, dPPR^*rbcL*^, made of consensus PPR motifs that were designed to bind a sequence near the 5’ end of *rbcL* transcripts in Arabidopsis chloroplasts. We used a functional complementation assay to demonstrate that this protein bound its intended RNA target with specificity *in vivo* and that it substituted for a natural PPR protein by stabilizing processed *rbcL* mRNA. We targeted a second protein of analogous design to the *petL* 5’ UTR, where it substituted for the native stabilizing PPR protein PGR3, albeit inefficiently. These results showed that artificial PPRs can be engineered to functionally mimic the class of native PPR proteins that serve as physical barriers against exoribonucleases.

## INTRODUCTION

The manipulation of gene expression represents a major challenge for both basic and applied biology. Progress in this field has been made possible by the discovery of natural products holding the potential to be tailored to powerful synthetic tools for genetic engineering. Posttranscriptional mechanisms play a prominent role in the control of gene expression and RNA binding proteins mediate these processes. Thus, the possibility to engineer RNA binding proteins with desired RNA binding specificity has attracted considerable attention (reviewed in 1, 2). Pentatricopeptide repeat (PPR) proteins constitute one of the largest families of RNA binding proteins in eukaryotes comprising more than 400 members in higher plants (3). PPR proteins are nucleus-encoded proteins but they function almost exclusively in mitochondria or chloroplasts where they hold various biological activities: protein barriers to RNA degradation, translational activation, recruitment of effectors to specific RNA sites, regulation of important RNA *cis*-element and remodeling of local RNA structures (reviewed in 4). PPR proteins are characterized by a variable number of tandem repeats (from 2 to over 30) of a degenerate 35 amino acid motif that forms a helix-loop-helix structure (5,6). The consecutive repeats stack to form a right-handed super helix that binds RNA with sequence specificity in a one-repeat/one-nucleotide mechanism (7). The RNA base bound by a PPR repeat is primarily determined by the identity of amino acids at two positions, 5 and 35 (8–11). The modular architecture of PPR repeats and the existence of an amino acid code for RNA base recognition make them an attractive scaffold for the rational design of RNA binding proteins with desired sequence specificity and therefore, the control of RNA functions *in vivo*. In fact, the PPR code has been successfully used to reprogram natural PPR proteins in plants to bind new mRNA sequences *in vivo* that are different from their native ones. For example, the PPR code was used to reprogram the sequence specificity and *in vivo* function of the mitochondrial PPR protein, RPF2 in Arabidopsis plants (12). RPF2 possesses 16 PPR repeats and targets two RNA sites sharing a strong sequence identity that are located within the 5’-UTRs of *cox3* and *nad9* genes to define the 5’ end processing of these transcripts by promoting a likely 5’-3’ endonucleolytic activity (13). Colas des Francs-Small et al. modified the amino acid composition of the RFP2 PPR tract to reprogram its *in vivo* specificity and bind a new RNA target within the mitochondrial *nad6* ORF which induced its subsequent cleavage. Despite its relative success, the assay highlighted a major *ab initio* limitation for the engineering of natural PPR proteins: the nucleotide specificity of only two of the 16 PPR motifs in RFP2 could be manipulated, which greatly restricted the choice of the RNA target to a sequence sharing high identity with RFP2’s native targets in mitochondria. The difficulty to freely reprogram the binding specificity of natural PPR proteins was additionally illustrated by a recent study that exploited the maize RNA stabilizer and translation enhancer, PPR10 and its cognate chloroplast *atpH* binding site to build an inducible switch for the expression of plastid transgenes in tobacco (14). In this study, a variant of PPR10 was successfully expressed from the tobacco nuclear genome to stimulate the expression of a chloroplast transgene whose mRNA stability and translation were under control of a modified version of the native PPR10 binding site. As for RFP2, however, the modification of PPR10 sequence specificity did not go further than 2 nucleotides.

Thus, in these two instances, the relative success of manipulating the specificity of natural PPR proteins *in vivo* is overshadowed by the inability to fully customize all of their PPR repeats to bind any chosen RNA sequence *in vivo*. The incapacity to recode the specificity of some PPR motifs in natural PPR proteins lies in their amino acid inconsistencies at positions that determine base specificity in relation to the PPR code and their unpredictable contribution to RNA binding (6,15). In order to circumvent the limitation of natural PPR proteins, several synthetic PPR scaffolds have been created (16–18). These artificial PPR proteins (called as well designer PPRs, dPPRs) derived from consensus PPR motifs that offer predictable and reliable sequence specificity *in vitro* (19). Several *in vivo* applications have been envisioned for dPPRs and each of these applications derived from two main functions that are naturally occupied by PPR proteins in organelles: the sequestration of RNA from interaction with other proteins or the targeting of effectors to specific RNA sites (reviewed in 20). Therefore, artificial PPR proteins must fulfill these two activities *in vivo* in order to be implementable as tools for the manipulation of RNA functions in living organisms. Currently, there is only one example for the *in vivo* application of dPPRs (21). In this study, a dPPR protein was successfully engineered in transgenic Arabidopsis plants to capture a specific mRNA in chloroplasts, demonstrating that the artificial dPPR scaffold can selectively and reliably bind designated RNA *in vivo*. However, this study did not reveal whether dPPRs hold functional potentialities similar to that of natural PPRs *in vivo*.

Many natural PPR proteins are involved in the control of RNA stability in plant organelles where they specifically bind to intergenic regions of RNA precursors to stabilize and protect the adjacent RNA sequences from degradation by 5’→3’ or 3’→5’ exoribonucleases therefore, defining the 5’ or 3’ ends of the processed RNAs. To test whether dPPRs can fulfill these properties *in vivo*, we conducted functional complementation assays in Arabidopsis of natural PPR proteins that stabilize the 5’ end of mature chloroplast mRNAs by an RNase blockade mechanism. We showed that a dPPR made of 13 PPR repeats and programmed to bind the *in vivo* 5’ end of a particular mRNA efficiently substituted for the biological function of the endogenous PPR protein. Furthermore, we demonstrated that positioning the RNA binding site of the dPPR to a different genomic location than that of the natural PPR protein creates a new and functional 5’ end of the processed mRNA *in vivo*. In conclusion, these results showed that dPPRs hold functional capacities similar to those of natural PPRs by sequestering specific RNA sequences and preventing their access to exoribonucleases. This study provides an additional application of artificial dPPRs in the targeted control of RNA stability *in vivo*.

## MATERIAL AND METHODS

### Plant material

The *mrl1* Arabidopsis T-DNA homozygote line (SALK_072806, Col-0 background) seeds were a generous donation of Olivier Vallon (22). Arabidopsis plants were grown on soil under controlled conditions: 12-h light (21°C): 12-h dark (18°C); 160-175 μmol photons m^-2^ sec^-1^. *Nicotiana benthamiana* plants were grown on soil under standard greenhouse conditions: 16-h light (22°C): 8-h dark (18°C) cycles; 200-250 μmol photons m^-2^ sec^-1^. Seeds for *pgr3-4* homozygote line (FLAG_086B06, Was-0 background) were a gift of Toshiharu Shikanai (Kyoto University) and is the same allele previously analyzed by ribosome profiling (23). Seeds were germinated on MS medium supplemented with 2% (w/v) sucrose and grown in at 22°C s in 10-h light: 14-h dark; 80 μmol photons m^-2^ sec^-1^ for 14 day before being transferred to soil.

### Complementation of *mrl1* and *pgr3* mutants with dPPR-encoding transgenes

The DNA sequences of Arabidopsis chloroplast transit peptide RecA and dPPR^*rbcL*^ were codon-optimized for Arabidopsis and synthesized by GeneCust company. The two sequences were assembled using overlapping PCRs. RecA and dPPR^*rbcL*^ were PCR-amplified using primers K450 Fw/K628 Rev and K627 Fw/K465 Rev, respectively, and assembled by overlapping PCR. Cloning was performed using Gateway technology (Thermo Fisher Scientific, Waltham, MA, USA) following the manufacturer’s protocols. The *RecA-dPPR^rbcL^* gene was first cloned into pDONR207 entry vector using BP clonase II and subsequently cloned into binary vectors pAUL1 (for C-terminal protein fusion with 3xHA tag) (24) or pMDC83 (for C-terminal protein fusion with GFP) (25) using LR clonase II. GV3101 Agrobacteria carrying the *RecA-dPPR^rbcL^*:pAUL1 construct were used for floral dip transformation of homozygous *mrl1* plants and Agrobacteria carrying the *RecA-dPPR^rbcL^*:pMDC83 construct were used for tobacco leaf agroinfiltration and protein subcellular localization. Seeds obtained from T_0_ Arabidopsis plants were sown on soil and the transgenic seedlings were selected for resistance to BASTA^®^ (Bayer AG, Leverkusen, Germany). 30 confirmed transgenic plants were screened for dPPR^*rbcL*^ expression by immunoblot on leaf total protein extracts using HA antibodies (H9658 clone, Sigma-Aldrich) and three lines expressing gradual levels of dPPR^*rbcL*^ protein were chosen for phenotypic analyses.

The dPPR^*petL*^ sequence was codon optimized for Arabidopsis, synthesized by Genewiz (South Plainfield, New Jersey) and cloned into a modified version of pCambia1300 that allows the C-terminal fusion of the protein with 3xFLAG, as described previously (21). This plasmid was used to transform Arabidopsis *pgr3-4* plants by the floral dip method. Transgenic seedlings were selected for resistance to hygromycin on Murashige and Skoog medium. Plants were grown for two weeks and pools of 6-to-8 seedlings were harvested for protein and RNA analysis.

Information about the primer sequences can be found in Supplementary Table 1. The DNA and protein sequences used in this study are provided in Supplementary Figure 1.

### Immunoblot analysis

Total protein extraction was achieved by homogenizing leaf discs (9 mm diameter) in 150 μl of 2x Laemmli sample buffer (120 mM Tris-HCl, pH 6.8, 4% SDS, 20% glycerol, 2,5% ß-mercaptoethanol, 0.01% bromophenol blue). Samples were centrifuged at 18,000 g for 5 min at room temperature and 10 μl of the supernatant were analyzed by SDS-polyacrylamide gel electrophoresis (10% polyacrylamide). Proteins were transferred onto PVDF membrane by wet western transfer using the Mini Trans-Blot^®^ Cell Assembly (Bio-Rad) in buffer containing 25 mM Tris, 192 mM glycine, 20% ethanol.

PsaD, AtpA, AtpB and PetD antibodies were described previously (26). The NdhL antibody was a gift of Toshiharu Shikanai (Kyoto University). PsbE and PsbA (D1) antibodies were purchased from Agrisera. Monoclonal anti-HA (H9658 clone), anti-FLAG (M2 clone) and anti-Myc (9E10 clone) antibodies were purchased from Sigma-Aldrich.

### Subcellular protein localization

*A. tumefaciens* strain GV3101 carrying the *RecA-dPPR^rbcL^*:pMDC83 construct were cultured overnight, pelleted for 5 min at 3,200 g and resuspended in 10 mM MES pH 5.6, 10 mM MgCl_2_, 150 μM acetosyringone to OD_600_ of 0.3. The bacterial suspension was incubated for 2 h at room temperature in the dark and used for infiltration of leaves of 3-week-old *N. benthamina* plants. Two days after infiltration, leaves were digested in protoplast extraction medium (0.01% Macerozyme R10, 0.1% Driselase, 0.2% Cellulase “Onozuka”, 4.3 g/L MS salt mix, 0.5 g/L MES pH 5.6, 20 g/L sucrose, 80 g/L mannitol) in the dark for 5 h at 30°C with gentle shaking (50 rpm). Protoplasts were examined under a Zeiss LSM 780 confocal microscope. GFP was excited with a 488 nm laser and emission was acquired between 493-556 nm. RFP and chlorophyll were excited with a 561 nm laser line and emissions were acquired between 588-641 nm and 671-754 nm, respectively.

### Purification of recombinant dPPR^*rbcL*^ protein

DNA encoding *dPPR^rbcL^* without the transit peptide sequence was PCR-amplified (K461 Fw/K462 Rev) and cloned into the pMAL-TEV vector using BamHI/SalI restriction sites. Rosetta 2 E. coli cells containing the protein expression vector were grown to OD_600_ of 0.5 and protein expression was induced with 1 mM IPTG for 3.5 h at 20°C and 220 rpm shaking. Bacteria were pelleted and resuspended in lysis buffer (30 mM Tris pH 7.5, 450 mM NaCl, 5 mM β-mercaptoethanol, complete™ EDTA-free Protease Inhibitor Cocktail (Roche) and lysed by sonication. The recombinant dPPR^*rbcL*^ fused to the maltose binding protein was affinity purified by binding to an amylose affinity column, washing three times with lysis buffer and eluting in the same buffer supplemented with 20 mM maltose. The recombinant MBP fusion protein was concentrated with an Amicon Ultra-15 centrifugal filter unit (Ultracel 3K; Merck Millipore, Darmstadt, Germany) and used directly for electrophoretic mobility shift assay (EMSA).

### Gel Mobility Shift assays

Synthetic RNA oligos were 5’ end-labeled with [gamma-^32^P]-ATP. Binding was performed for 3 h at 25°C in 20 μl reactions containing increasing amounts of rdPPR^*rbcL*^, 20 pM of RNA, 100 mM NaCl, 40 mM Tris pH 7.5, 4 mM DTT, 0,5 mg/ml Heparin, 10% glycerol, and 1 U RNasin Plus RNase Inhibitor (Promega). Reactions were separated on native 5% polyacrylamide gels in 0.5x TBE buffer at 4°C and gels were dried after electrophoresis and exposed to a phoshorimaging plate.

### *In vivo* protein labeling

*In vivo* labeling was performed on leaf discs from 4-week-old plants as described (27). Labeling was carried out for 15 min and total proteins were extracted as described previously.

### RNA gel blot hybridization, cRT-PCR and primer extension assays

Total RNA was extracted from 2-week-old plants using Tri-Reagent (Molecular Research Center, Inc.) according to the TRIzol protocol (Thermo Fisher Scientific, Waltham, MA, USA). RNA was further purified by an additional phenol/chloroform extraction and ethanol precipitation. Five micrograms of total leaf RNA were resolved on denaturing formaldehyde gels as desribed (28). RNA was blotted onto Hybond-N^+^ membrane (GE Healthcare) by capillary transfer overnight using 20x SSC as blotting buffer. 60-mer DNA oligonucletoides were 5’ end-labeled using [γ-^32^P]-ATP and PNK (Thermo Fisher Scientific, Waltham, MA, USA) according to the manufacturer’s instructions. Blots were hybridized overnight at 50°C in Church buffer (7% SDS, 0.5 M NaPhosphate pH 7.0, 1 mM EDTA) and washed twice with washing buffer (1x SSC, 0,1% SDS) for 5 minutes at the hybridization temperature. Results were visualized on an Amersham Typhoon imager. Analysis of *petL* RNA by RNA gel blot hybridization used 6 μg leaf RNA and a 5’-end labeled synthetic DNA probe complementary to the *petL* open reading frame. The blots were hybridized overnight at 48°C in Church buffer and washed five times for 10 min in 0.2% SDS and 5X SSC at 48°C.

cRT-PCR was carried out as described (29) except that SSIV RT (Thermo Fisher Scientific, Waltham, MA, USA) and GoTaq (Promega) enzymes were used. K801 Rev primer was used for cDNA synthesis and K801 Rev/K803 Fw primers for the subsequent PCR. To identify the primary transcript ends, RNA was treated with RNA 5’ Pyrophosphohydrolase (NEB) according to the manufacturer’s instructions prior to RNA circularization. cRT-PCR products were cloned into pGEM^®^-T Easy vector (Promega) and more than 10 clones/cRT-PCR product were sequenced.

For primer extension assays of *rbcL*, 10 μg of total RNA were incubated in the presence of [gamma-^32^P]-ATP 5’ end-labeled K626 Rev primer and 0.5 mM dNTPs at 75°C for 5 min. Temperature was then reduced to 50°C and SSIV RT enzyme mix (Thermo Fisher Scientific, Waltham, MA, USA) was added according to the manufacturer’s protocol for First-Strand cDNA Synthesis Reaction. Reactions were incubated for 30 min at 50°C and stopped by adding one volume of RNA loading buffer (90% deionized formamide, 20 mM Tris pH 8.0, 20 mM EDTA, traces of bromophenol blue and xylene cyanole) and heating for 5 min at 95°C. Extension products were loaded onto an 8% polyacrylamide gel containing 8 M urea and run in 1x TBE buffer. The DynaMarker^®^ Prestain Marker for Small RNA Plus (BioDynamics Laboratory Inc.) was run in parallel on the gel.

Primer extension analysis of the *petL* 5’ end was performed with leaf RNA (2 μg of the wild-type and 7 μg for the *pgr3* mutant) and a 5’-end labeled synthetic DNA oligonucleotide as described previously (30). Products were resolved in an 8% denaturing polyacrylamide gel, and imaged with a Storm phosphorimager.

### RIP-Seq and slot blot analysis

RIP-Seq analysis was performed as described previously (31). Briefly, stromal extracts (1 mg protein) were isolated from 2-week-old dPPR^*rbcL*^-1 or Col-0 plants and incubated with monoclonal anti-HA antibodies (Sigma). IgGs were captured with Protein A DynaBeads (Thermo Fisher Scientific, Waltham, MA, USA) and co-precipitated RNA was recovered by Trizol extraction followed by phenol/chloroform extraction and ethanol precipitation. The RNA was purified with the Monarch^®^ RNA Cleanup Kit (NEB). 50 ng of RNA from each experiment were used for library generation with the NEBNext^®^ Ultra™ II RNA Library Prep Kit according to the manufacturer’s instructions. Deep sequencing (2x 250 bp, v3 chemistry) was performed on a MiSeq sequencer (Illumina, San Diego, CA, USA) yielding 13.0 and 14.7 Mio (dPPR^*rbcL*^-1 replicate 1 and 2, respectively) as well as 15.5 and 13.1 Mio (Col-0 replicate 1 and 2, respectively) trimmed paired reads. To determine depth of coverage (reads/nucleotides), the primary reads were aligned to the Arabidopsis chloroplast genome (accession number NC_000932.1) using CLC Genomics Workbench 6.5.1 (Qiagen, Valencia, CA, USA) with the following parameters: *mismatch cost = 2, insertion cost = 3, deletion cost = 3, length fraction = 0.5, similarity fraction = 0.8, global alignment = no, auto-detect paired distances = yes*. Aligned reads were extracted as reads per nucleotide and the mean value of the two replicates was displayed across the entire chloroplast genome.

For slot blot analysis, the co-immunoprecipitated RNA was recovered as described above. 1/4^th^ of the pellet RNA and 1/40^th^ of the supernatants were heated at 70°C for 10 min in 2x SSPE buffer (300 mM NaCl, 20 mM NaH2PO4, and 2 mM EDTA at pH 7.4) and blotted onto Hybond-N^+^ membrane (GE Healthcare) using a Minifold I slot blot system (GE Healthcare). Probe labeling and hybridization were as described for RNA gel blot analysis. Some of the blots were stripped to be rehybridized with another gene probe. Results were visualized on an Amersham Typhoon imager and data quantification was performed with ImageQuant TL (GE Healthcare).

## RESULTS

### Customization of a dPPR targeting the 5’ end of mature *rbcL* mRNA

To test whether artificial PPRs built from consensus PPR motifs have similar functional capacities as natural P-type PPRs *in vivo*, we used an *in vivo* functional complementation assay of the Arabidopsis PPR protein MRL1. MRL1 is a PPR protein that is targeted to chloroplasts, where it stabilizes a processed isoform of RNA from the *rbcL* gene. *rbcL* encodes the large subunit of Rubisco and is transcribed into a primary transcript whose 5’ end maps at position −177 from the start codon (Figure 1A) (22,32,33). This mRNA precursor is processed to a transcript possessing a shorter 5’-UTR mapping at position −69. Accumulation of the processed isoform requires MRL1, and processing is presumed to involve the canonical PPR barrier mechanism: 5’-to-3’ exonucleolytic degradation back to the MRL1 barrier bound immediately downstream of −69 (22). The RNA fragments bound by PPR proteins that stabilize RNA usually accumulate as small RNA footprints (sRNAs) whose termini coincide with the positions of the 5’ or 3’ ends of the mRNAs they stabilize *in vivo* (34–36). The MRL1 RNA footprint accumulates as a ~20-30 nt sRNA whose 5’-end coincides with the position of the processed *rbcL* mRNA, due to protection by the bound protein (Figure 1A). Despite the loss of the processed *rbcL* mRNA, *mrl1* mutant plants do not show a particular growth or physiological defect besides a slight reduction of RbcL protein content, suggesting that the primary *rbcL* transcript is translationally competent to produce sufficient RbcL (22). Therefore, *mrl1* serves as an ideal surrogate plant to express an artificial PPR and test its ability to complement the *in vivo* function of a natural PPR in gene-specific RNA stabilization by monitoring the recovery of the mature *rbcL* mRNA.

**Figure 1.**
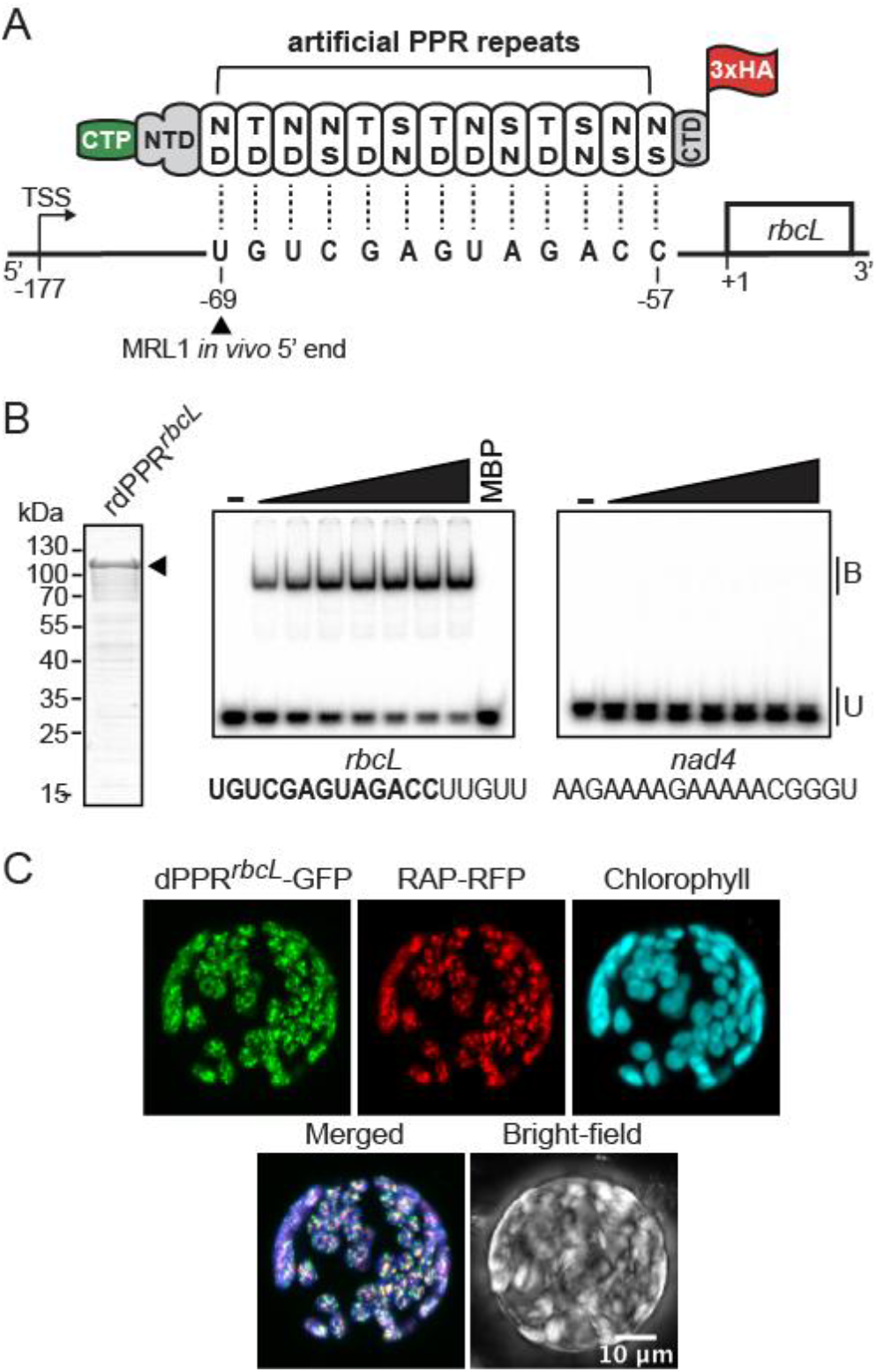
Engineering of an artificial PPR protein (dPPR^*rbcL*^) programmed to bind the 5’ end of mature *rbcL* mRNA in Arabidopsis chloroplasts. (**A**) Schematic of dPPR^*rbcL*^ design. The engineered protein was made of 13 consensus PPR tandem repeats programmed to bind a 13 nucleotide sequence beginning 69 nt and finishing 57 nt upstream of *rbcL* start codon. The dPPR tract was flanked by the N- and C-terminal domains (NTD and CTD) of the maize PPR10 protein. The top and bottom amino-acids featured in each PPR repeat correspond to the amino acid positions 5 and 35 that determine base specificity. An N-terminal chloroplast transit peptide from RecA protein (CTP) and a C-terminal 3xHA epitope tag were added to the protein to allow the chloroplast import and immunodetection of the protein *in vivo*. The *rbcL* transcription start site maps at position −177 and the wild-type (WT) *in vivo* 5’ end of processed *rbcL* mRNA at position −69. The MRL1 footprint is underlined. The genomic positions are given according to *rbcL* start codon. (**B**) Electrophoretic mobility shift assay (EMSA) showing preferential binding of recombinant dPPR^*rbcL*^ (rdPPR^*rbcL*^) to *rbcL* 5’ end. *(Left)* An aliquot of the purified rdPPR^*rbcL*^ used for binding assays was analyzed by SDS-PAGE and staining with Coomassie Blue. rdPPR^*rbcL*^ run on the gel at a predicted size of ~118 kDa and is indicated by an arrowhead. (*Right*) rdPPR^*rbcL*^ was used in EMSAs with radiolabeled RNA oligonucleotides (*rbcL* or *nad4*) whose sequences are shown below. The sequence of the designated dPPR^*rbcL*^ binding site is highlighted in bold. Protein concentrations were 22, 44, 66, 88, 110, and 132 nM for rdPPR^*rbcL*^ and 200 nM for the maltose binding protein (MBP). Bound (B) and unbound (U) RNAs are indicated. (**C**) Chloroplast localization of dPPR^*rbcL*^ in *N. benthamiana* cells. The dPPR^*rbcL*^-GFP fusion protein was transiently co-expressed in tobacco protoplasts with the Arabidopsis chloroplast nucleoid-associated protein RAP fused to RFP. Fluorescence images of GFP, RFP, chlorophyll and a merged image are shown.

To this end, we conceived a protein, dPPR^*rbcL*^, targeting the *in vivo* 5’-end of processed *rbcL* mRNA according to the artificial PPR design described by Shen et al. (16). The dPPR^*rbcL*^ was made of 13 consensus PPR repeats flanked by amino- and carboxy-terminal segments of the maize chloroplast-localized protein PPR10 (30), and the PPR tract was programmed to bind a 13 nucleotide sequence matching the processed 5’ end of the mature *rbcL* mRNA (Figure 1A). We chose a 13-motif design for dPPR^*rbcL*^ due to evidence that longer artificial PPR tracts are prone to increased off-target binding (19). For the *in vivo* complementation assay, the dPPR^*rbcL*^ additionally included the N-terminal chloroplast transit peptide of Arabidopsis RecA (first 68 amino acids) whose efficiency to target exogenous proteins to chloroplasts had already been demonstrated (37) and a C-terminal 3xHA tag for immunodetection (Figure 1A and Supplementary Figure 1). As a preliminary step to the functional complementation assay *in vivo*, we confirmed the RNA sequence binding specificity of dPPR^*rbcL*^ by expressing and purifying the mature recombinant protein (lacking the cleaved transit peptide), rdPPR^*rbcL*^ in *E. coli* as an N-terminal protein fusion with the maltose binding protein and we used this protein in gel mobility shift assays (Figure 1B). The rdPPR^*rbcL*^ bound a 18-nt RNA matching the *in vivo* 5’ end sequence of processed *rbcL* mRNA but did not bind an unrelated mitochondrial RNA sequence of the same size (*nad4*) demonstrating that, as expected, dPPR^*rbcL*^ binds with specificity to its designated RNA target *in vitro*. Furthermore, rdPPR^*rbcL*^ markedly bound its RNA ligand at protein concentrations in the nanomolar range showing that it binds RNA with high affinity.

Next, we tested the subcellular localization of the chimeric dPPR^*rbcL*^ protein in plant cells. To this end, a dPPR^*rbcL*^-GFP fusion protein was transiently expressed in *Nicotiana benthamiana* leaves and leaf protoplasts were examined by confocal microscopy (Figure 1C). The GFP signal colocalized with the chlorophyll fluorescence demonstrating that dPPR^*rbcL*^-GFP was efficiently imported to chloroplasts. In addition, the protein was found in discrete foci within chloroplasts. These foci colocalized with the fluorescence signal of a co-expressed chloroplast protein RAP fused to RFP that was previously shown to associate with the chloroplast nucleoids *in vivo* (38), indicating that dPPR^*rbcL*^ associates with the nucleoids as well. This observation is in agreement with the reported association of MRL1 and several other PPRs controlling mRNA stability with the nucleoids of chloroplasts in maize (39). The results demonstrated that the customized dPPR^*rbcL*^ was expressed in plant cells where it localized to the chloroplasts similarly to its natural PPR counterpart, MRL1.

### dPPR^*rbcL*^ can substitute for MRL1 to stabilize the 5’ end of *rbcL* mRNA *in vivo*

To test the *in vivo* activity of dPPR^*rbcL*^ and its capacity to complement the biological function of MRL1, we generated transgenic Arabidopsis plants expressing dPPR^*rbcL*^ in the *mrl1* mutant background. Three independent transgenic lines expressing different levels of dPPR^*rbcL*^ protein (Figure 2A) were selected for further phenotypic analyses. The expression of dPPR^*rbcL*^ in the transgenic plants did not cause any growth phenotype as compared to *mrl1* and wild-type (Col-0) plants (Figure 2B), and immunoblot analyses using antibodies against core subunits of the major thylakoid membrane complexes did not reveal any change in chloroplast protein accumulation (Figure 2A). These results suggest that expression of dPPR^*rbcL*^ does not cause deleterious pleiotropic effects in plants.

**Figure 2.**
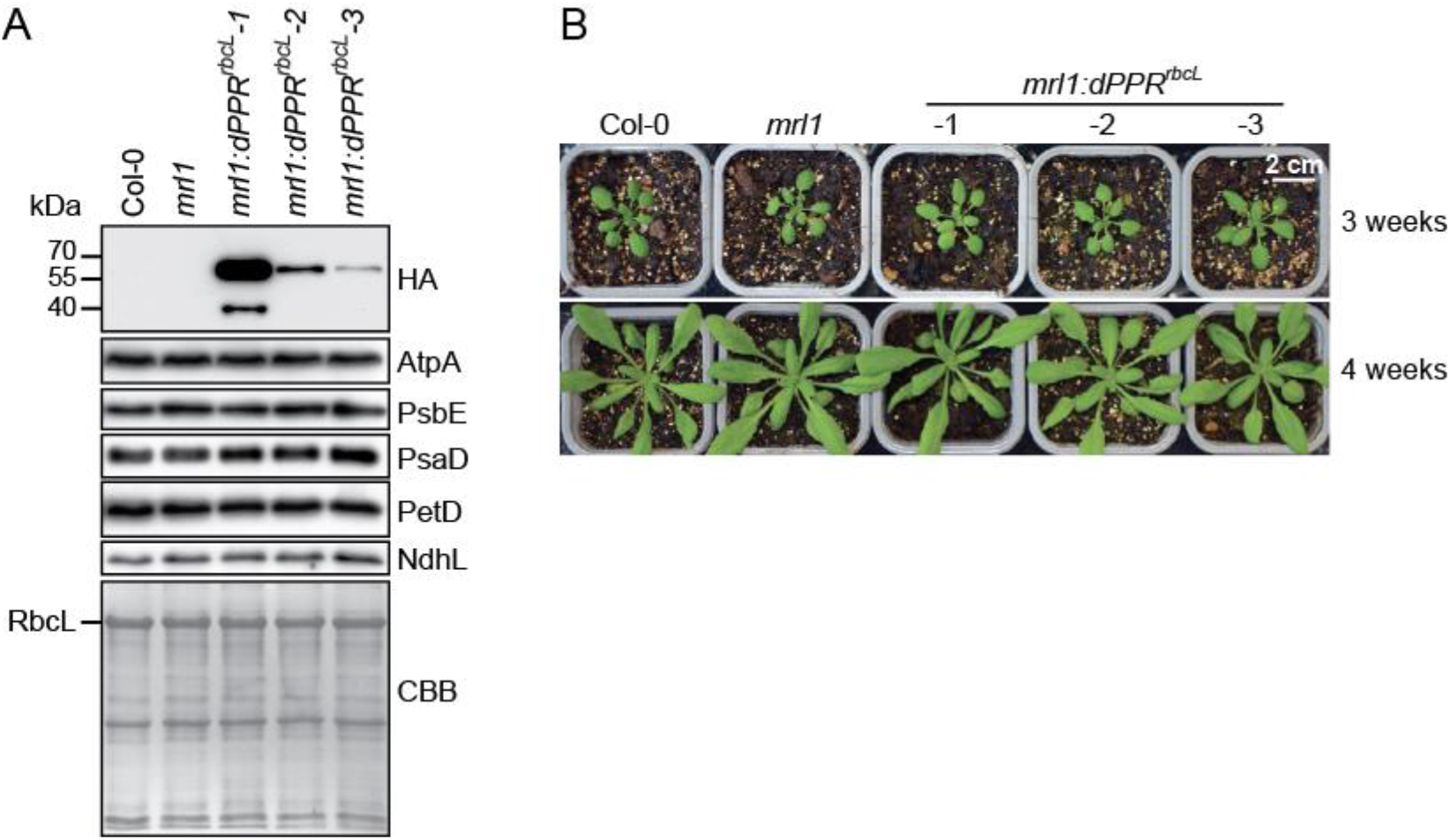
Phenotype of transgenic *mrl1* Arabidopsis plants expressing dPPR^*rbcL*^. (**A**) Three *mrl1:dPPR^rbcL^* independent lines expressing gradual levels of dPPR^*rbcL*^ were analyzed by immunoblots on total leaf protein extracts along with wild type (Col-0) and *mrl1* mutant plants. The dPPR^*rbcL*^ was detected with antibodies against the HA tag. The abundance of the chloroplast photosynthetic enzyme complexes in the different plant genotypes were analyzed with antibodies against core subunits: AtpA (ATP synthase), PsbE (Photosystem II), PsaD (Photosystem I), PetD (Cytochrome *b_6_f*) and NdhL (NADH dehydrogenase). One of the membrane was stained with Coomassie brilliant blue (CBB) serve as the protein loading control. The band corresponding to the large subunit of Rubisco (RbcL) is marked. (**B**) Visible phenotype of 3- (top) and 4-week-old (bottom) Col-0, *mrl1* and transgenic *mrl1:dPPR^rbcL^* Arabidopsis plants.

To determine whether dPPR^*rbcL*^ stabilized the 5’ end of processed *rbcL* mRNAs in the *mrl1* mutant, we performed a primer extension assay that allows the identification of both the primary and 5’-end processed *rbcL* transcripts (Figure 3A). As expected, both the primary and processed *rbcL* mRNAs were detected in wild-type plants whereas the processed RNA was absent in *mrl1* mutant. However, we observed that *mrl1* mutant also showed some reduction in the primary *rbcL* mRNA suggesting that the protein might have a secondary binding site upstream, or that its absence might expose an otherwise inaccessible endonuclease-sensitive site. In the three transgenic *mrl1* lines expressing dPPR^*rbcL*^ protein, a processed *rbcL* mRNA isoform accumulated, and the abundance of this isoform correlated with the abundance of the dPPR^*rbcL*^ protein in each line. Altogether, these results demonstrated that the artificial dPPR^*rbcL*^ had the capacity to control the stability of its RNA target *in vivo* and to complement the function of the natural PPR MRL1.

**Figure 3.**
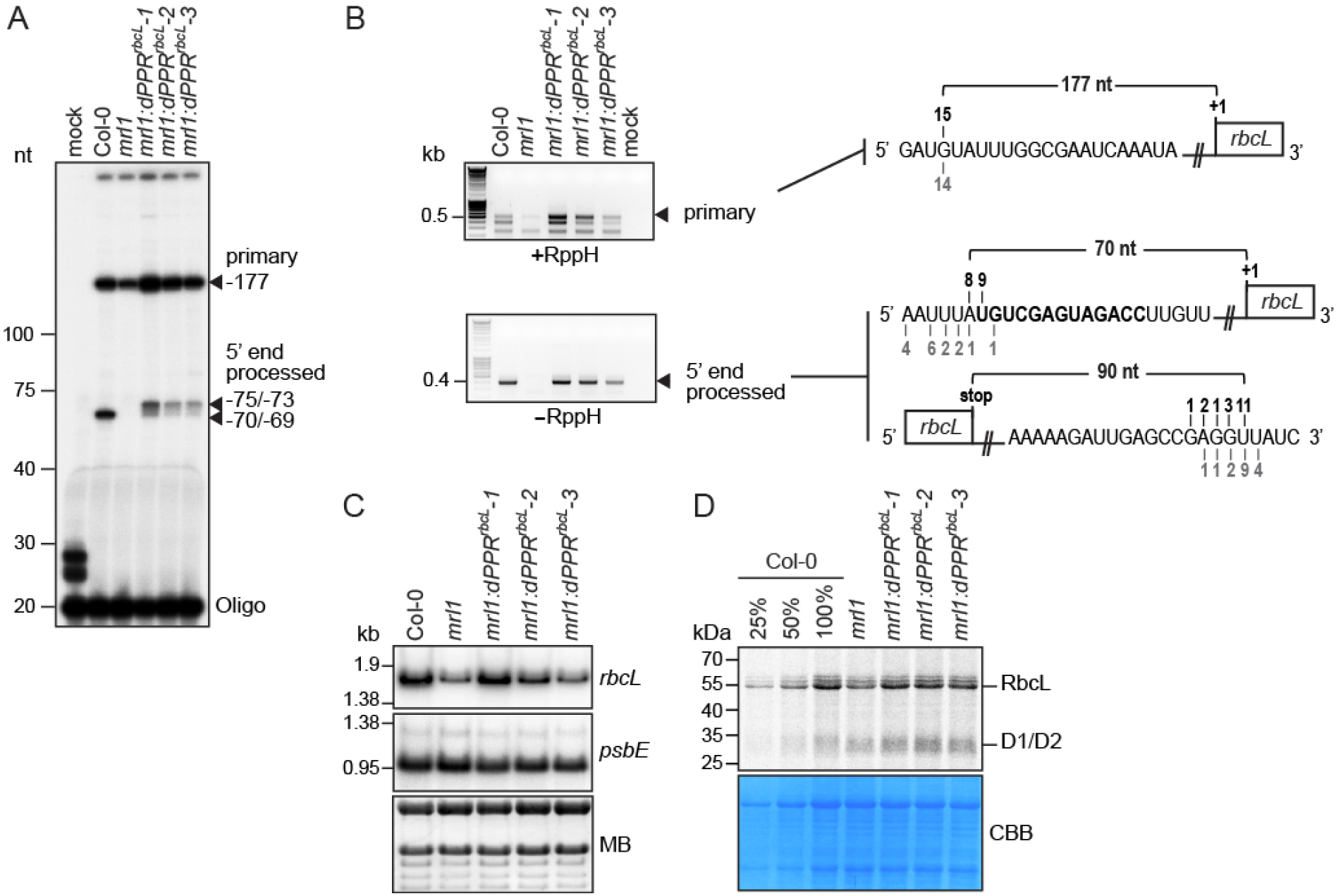
Functional complementation assay of *mrl1* Arabidopsis mutant by dPPR^*rbcL*^. (**A**) Primer extension analysis of *rbcL* mRNAs 5’ end accumulating in Arabidopsis chloroplasts. The extension products were separated on a 8% denaturing polyacrylamide gel. RNA samples from Col-0, *mrl1* and the transgenic plant lines were analyzed. The arrowheads indicate the 5’ end of the primarily transcribed and processed *rbcL* mRNA. Their positions are given according to the *rbcL* start codon and their mapping by cRT-PCR. (**B**) cRT-PCR mapping of the termini of the primary (+RppH) and processed (-RppH) *rbcL* mRNAs in the Col-0 and complemented *mrl1* plants. The arrowheads indicate the cRT-PCR bands that were excised from the agarose gel for sequencing. The diagrams on the left display the RNA sequences annotated with the 5’ or 3’ ends determined by cRT-PCR. The dPPR^*rbcL*^ binding site is highlighted in bold and the number of clones mapping to each position is indicated above and underneath the RNA sequence for the Col-0 and *mrl1:dPPR^rbcL^* genotypes, respectively. RppH: Pyrophosphohydrolase. (**C**) RNA gel blot hybridizations of leaf RNAs from plants of the indicated genotypes using strand-specific oligonucleotide probes for *rbcL* and a chloroplast control *psbE* gene. A portion of one of the blots stained with methylene blue (MB) is shown to illustrate equal sample loading. (**D**) *In vivo* chloroplast protein synthesis analysis. Intact leaves from the indicated plant genotypes were ^35^S pulse labelled in presence of cycloheximide to block cytosolic translation and ^35^S-labeled leaf proteins were fractionated by SDS-PAGE. The autoradiograph and Coomassie blue stained gel (CBB) pictures are shown. The signal bands corresponding to synthesized RbcL and D1/D2 chloroplast proteins are indicated.

The primer extension analysis revealed that the processed 5’ end of *rbcL* mRNA that accumulated in the transgenic plants was several nucleotides longer than that in the wild-type plants. We conducted circular RT-PCR to map precisely the ends of the different *rbcL* mRNAs accumulating in the different genotypes (Figure 2B). An RNA treatment with the 5’ pyrophosphohydrolase prior to cRT-PCR allowed to distinguish the 5’ ends of the primary *rbcL* mRNA from the processed ones. In agreement with the primer extension results, the cRT-PCR analysis revealed that the 5’ end of the primary transcript mapped 177 nt upstream of the *rbcL* start codon in all genotypes. In addition, the results confirmed that the major 5’ ends of the processed *rbcL* mRNA in the transgenic plants mapped at position −75/−73 from the *rbcL* start codon, 4 to 6 nucleotides upstream of the processed 5’ end in the wild-type. The position of the 5’-end that is stabilized by dPPR^*rbcL*^ is as expected based on the fact that PPR10 (from which the N-terminal region of this dPPR is derived) protects several nucleotides upstream of the sequences bound by its canonical PPR motifs (40). Since we designated the first PPR repeat of dPPR^*rbcL*^ to bind the immediate 5’ nucleotide of the MRL1-dependent processed *rbcL* mRNA, the engineered protein was expected to stabilize a processed *rbcL* mRNA with an end mapping several nucleotides 5’ to that in the wild-type.

Consistent with the loss of the mature *rbcL* mRNA in *mrl1* mutant, the examination of the accumulation of *rbcL* transcripts by RNA gel blot hybridization showed that the level of *rbcL* mRNAs was reduced in the mutant compared to the wild-type (Figure 3C). In the complemented *mrl1* lines, *rbcL* mRNA level was restored according to the level of dPPR expression in these plants with a full recovery of mRNA abundance in the strong expressor line that was comparable to the wild-type level.

Despite the reduction of *rbcL* mRNA abundance in *mrl1*, Coomassie staining of total leaf protein resolved by SDS-PAGE did not reveal a noticeable reduction of RbcL protein in the mutant compared to the wild type (Figure 2A). However, the Coomassie staining might not be sufficiently sensitive to detect subtle changes of RbcL content. Therefore, we measured the accumulation of newly synthesized RbcL in the different genotypes by pulse labeling analysis using intact plant leaf tissues fed with the ^35^S-radiolabeled methionine (Figure 3D). The results showed that the loss of *rbcL* mRNA in *mrl1* was accompanied by a moderate reduction in the rate of RbcL synthesis. The level of *rbcL* mRNA correlated well with the abundance of neosynthesized RbcL protein in transgenic, wild-type and *mrl1* mutant plants. These results indicate that the 5’ end processed *rbcL* mRNA contributes to the accumulation of RbcL protein in plants and confirmed that the 5’ end processed *rbcL* mRNA that was newly defined by dPPR^*rbcL*^ in chloroplasts is functional in the transgenic plants.

Altogether, the molecular analyses demonstrated that the artificial dPPR^*rbcL*^ protein has the capacity to complement the *vivo* function of the PPR protein, MRL1 and control RNA stability in chloroplasts.

### dPPR^*rbcL*^ binds preferentially to the 5’ UTR of *rbcL* mRNA *in vivo*

To get a genome-wide view of dPPR^*rbcL*^ RNA binding specificity *in vivo*, we performed RIP-seq analysis using chloroplast stroma isolated from the *dPPR^rbcL^* transgenic and wild-type plants in two replicate experiments (Figure 4A). The read values (reads/nucleotide) and read mapping of the two immunoprecipitation replicates for each genotype (wild type and *mrl1:dPPR^rbcL^*) were highly reproducible (Supplemental Figure 2). Therefore, the mean read values of the two replicates from the experimental and control immunoprecipitations were aligned to the Arabidopsis chloroplast genome (Figure 4B). The RNA immunoprecipitation using wild-type stroma yielded very low read coverage of the chloroplast genome except for the region covering the rRNA genes (Figure 4B, Supplementary Figure 2) whereas the experiment using the transgenic plants yielded several local peaks in *psbA, atpF, psbC, psaB, rbcL, psbE* and *psbT* genes with *rbcL* peak being the most prominent (Figure 4B and Supplemental Table 2). Consistent with our experimental design, the genomic position of the prominent *rbcL* peak mapped ~30 nt upstream of the dPPR’s designated binding site (Supplemental Table 2, Supplementary Figure 3 and Figure 4C). Calculation of the enrichment of RNAs in the experimental versus control immunoprecipitations showed that RNAs in the 5’ UTR of *rbcL* were enriched more than 700-fold whereas RNAs mapping within the *rbcL* ORF were more weakly enriched (less than 100-fold) demonstrating that dPPR^*rbcL*^ primarily binds the 5’ UTR (Figure 4C). The RNAs from the other loci (*psbA, atpF, psbC, psaB, psbE* and *psbT*) were considered as potential off-targets and were examined in an independent RIP experiment. In this experiment, the experimental and control immunoprecipitations were both performed on stroma from the *mrl1:dPPR^rbcL^* transgenic plants but used different antibodies. The immunoprecipitated RNAs with HA antibodies were compared to those from an immunoprecipitation with Myc antibodies that do not recognize dPPR^*rbcL*^ protein and the recovered RNAs from the pellets and supernatants were subsequently analyzed by slot blot hybridizations (Figure 4D). In agreement with the RIP-seq analysis, the slot blot data confirmed that *rbcL, psbA, atpF, psbC, psaB, psbE, psbT* RNAs were specifically recovered in the dPPR^*rbcL*^ immunoprecipitation but with different degrees of enrichment. Quantification of the signals in the pellet versus supernatant showed that *rbcL* RNA was strongly enriched in the pellet whereas RNAs from *psbA, atpF, psbC, psaB, psbE* and *psbT* loci weakly coimmunoprecipitated with dPPR^*rbcL*^ (Figure 4D). Altogether, the data argue that dPPR^*rbcL*^ strongly associates to its designated *rbcL* RNA target *in vivo* while binding to a few off-targets with much less affinity.

**Figure 4.**
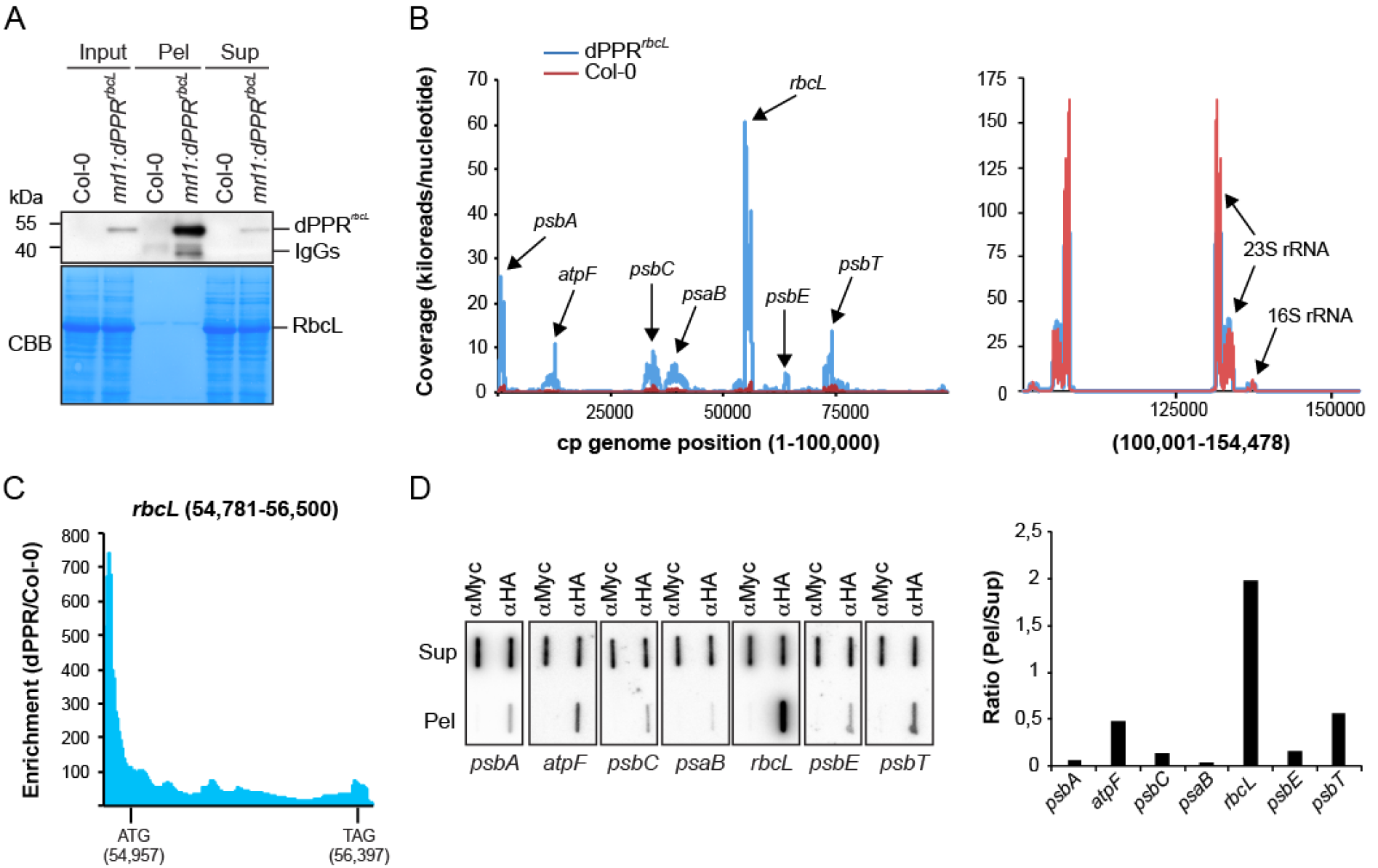
Chloroplast genome-wide analysis of RNAs associated with dPPR^*rbcL*^ *in vivo*. (**A**) Immunoblot analysis of an immunoprecipitation of dPPR^*rbcL*^ performed with HA antibodies on stroma extracts from the complemented *mrl1:dPPR^rbcL^* or Col-0 Arabidopsis plants (negative control). Pel: immunoprecipitation pellet, Sup: supernatant; IgG: Immunoglobuline G. IgGs in the experimental pellet are detected by the secondary antibody used to probe the immunoblot. A portion of the blot stained with Coomassie blue is shown to display equal loading. (**B**) RIP-seq analysis of chloroplast RNAs that associates with dPPR^*rbcL*^ *in vivo*. The mean coverage in kiloreads per nucleotide of two replicates for the experimental dPPR^*rbcL*^ immunoprecipitation and negative Col-0 control are plotted along the chloroplast genome on the same graph. Since the read coverage was too high for the chloroplast rRNA loci, these were split and displayed in a separate graph on the right. The main peaks are labeled with the names of the locus they belong to. Data for replicate experiments are shown in Supplementary Figure 2 and the read counts are provided in Supplementary Table 2. (**C**) Local enrichment (Ratio dPPR^*rbcL*^/Col-0) of *rbcL* RNA. The highest enrichment (more than 700-fold) is found in the 5’ UTR region where the binding site of dPPR^*rbcL*^ is located. The positions of the *rbcL* initiation (ATG) and stop (TAG) codons are indicated with their genomic position. (**D**) Slot blot hybridization analysis of RNAs that coimmunoprecipitate with dPPR^*rbcL*^ *in vivo*. The experimental and control immunoprecipitations were both performed with *mrl1:dPPR^rbcL^* stroma but used different antibodies: α-HA that detects dPPR^*rbcL*^ or α-Myc as a negative control. Four replicate blots were hybridized with strand and gene specific oligonucleotide probes. The same blot was used for hybridization with *psbT* and *psbE* probes, *psbC* and *psaB* probes and *psbA* and *atpF* probes after stripping. The hybridization signals in the pellet and supernatant of dPPR^*rbcL*^ immunoprecipitation were quantified for each gene and their ratio is displayed to the right.

### dPPR^*petL*^ can partially substitute for PGR3 to stabilize the 5’ end of *petL* mRNA *in vivo*

To provide a second example, we addressed whether an artificial PPR protein can functionally substitute for the Arabidopsis PPR protein PGR3. PGR3 binds the *petL* 5’-UTR, where it protects the downstream RNA from degradation and activates *petL* translation (41–43). PGR3 also binds the *rpl14-rps8* intergenic region, where it stabilizes a processed 3’-end and mildly stimulates *rps8* translation (23). An effect of PGR3 on NDH abundance has also been reported, but the basis for that effect remains unclear (23,41–43). Similarly to MRL1, the RNA footprint of PGR3 is represented *in vivo* by an abundant sRNA whose sequence matches the PGR3-dependent processed 5’-end of *petL* mRNA (34–36) (Figure 5A).

**Figure 5.**
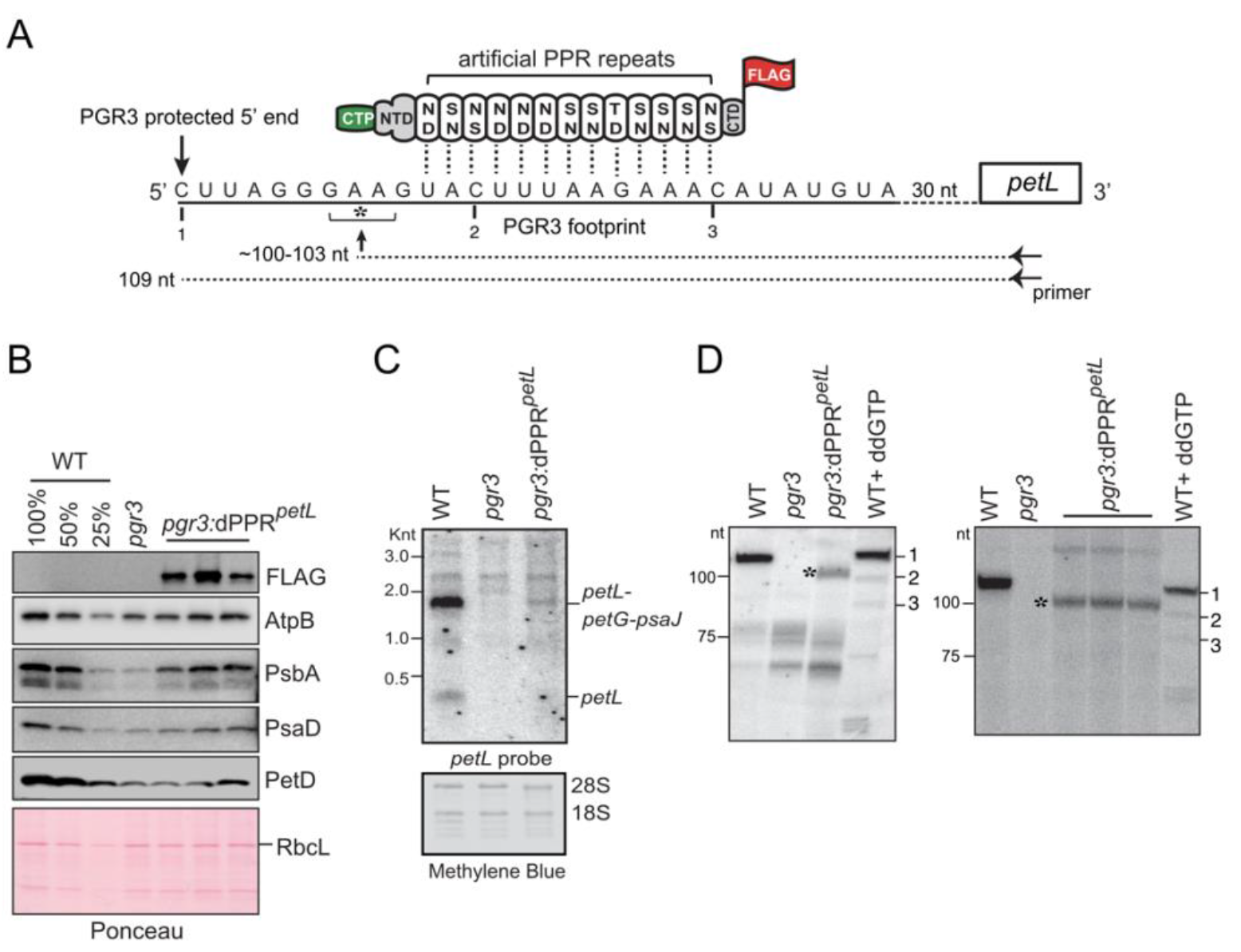
Partial complementation of PGR3’s *petL* RNA stabilization function by dPPR^*petL*^. **(A)** Schematic of dPPRdesign. The protein consists of 13 consensus PPR motifs programmed to bind the indicated sequence in the *petL* 5’UTR. The amino acids at positions 5 (top) and 35 (bottom) in each dPPR motif are indicated. The dPPR tract was flanked by the N- and C-terminal regions (NTD and CTD) of maize PPR10, including PPRIO’s chloroplast targeting peptide (CTP). A 3xFLAG tag was fused to the C-terminus. The 5’ end that is stabilized by PGR3 *in vivo* is marked with a vertical arrow, and the PGR3 footprint is underlined with a solid line. The horizontal arrow denotes the primer used for primer extension and the asterisk marks the 5’-end that is stabilized by dPPR’^*et*^. The nucleotides marked 1, 2, and 3 are primer extension stops resulting from the use of the chain terminator ddG, which serve as size markers on the primer extension gels in panel D. **(B)** Immunoblot analysis *pgr3* mutants expressing dPPR^*petL*^. Protein analyzed in each lane comes from pooled plants that were confirmed to have the indicated genotype by PCR. Replicate blots were probed with antibodies to the indicated proteins. The FLAG antibody detects dPPR^*petL*^. The Ponceau S-stained blot is shown below to illustrate equal sample loading and the abundance of the large subunit of Rubisco (RbcL). **(C)** RNA gel blot hybridization demonstrating partial rescue of *petL* transcripts in *pgr3* mutants expressing dPPR^*petL*^. 2 μg of leaf RNA was loaded in each lane. The methylene blue stained blot is shown below to demonstrate equal sample loading. 28S and 18S are cytosolic rRNAs. **(D)** Primer extension assays demonstrating stabilization of the expected novel *petL* 5’ terminus by dPPR^*petL*^. The two panels show analysis of four independent pools of *pgr3* mutants expressing dPPR^*petL*^. The positions of size markers are shown to the left. The last lane in each gel (WT + ddGTP) shows a primer extension reaction that used wild-type RNA and a trace amount of ddGTP to induce termination at C residues in the template (see panel A). The gel on the left used 4 μg of RNA from the WT (Ws-0) and *pgr3* samples, and 14 μg from *pgr3:dPPR^petL^*. The gel on the right used 2.5 μg of RNA from the WT (Ws-0) and *pgr3* samples, and 7.5 μg from *pgr3*:dPPR^*petL*^.

We designed an artificial PPR protein, dPPR^*petL*^ to bind a sequence within the PGR3 binding site in *petL*, but beginning several nucleotides downstream of the PGR3-dependent 5’ end (Figure 5A). The protein design is analogous to that of dPPR^*rbcL*^ except for the use of a different chloroplast targeting sequence (PPR10 versus RecA) and C-terminal tag (FLAG *versus* HA) (Supplementary Figure 1). The native PPR10 chloroplast targeting peptide was previously shown to target a dPPR to Arabidopsis chloroplasts *in vivo* (21). We introduced the transgene expressing dPPR^*petL*^ into the genome of a null *pgr3* mutant, and used immunoblot analysis to identify lines expressing dPPR^*petL*^ (Figure 5B). The *pgr3* mutant has a barely discernable phenotype under the growth conditions we used, and the expression of dPPR^*petL*^ did not have an obvious effect on the phenotype (Supplementary Figure 4). The *pgr3* mutant exhibited a modest decrease in subunits of the ATP synthase (AtpB), PSII (PsbA), PSI (PsaD), and the cytochrome *b6f* complex (PetD) (Figure 5B), consistent with the mild defect in chloroplast translation reported previously for the same null allele (23). Expression of dPPR^*petL*^ caused a slight increase in the abundance of PsbA and possibly several of the other proteins (Figure 5B). RNA gel blot hybridization showed the expected loss of *petL* transcripts from the *pgr3* mutant (Figure 5C). Expression of dPPR^*petL*^ resulted in partial restoration of the major PGR3-dependent transcript isoform (a tricistronic *petL-petG-psaJ* RNA) in the transgenic plants.

To probe further into the effects of dPPR^*petL*^ on *petL* RNA metabolism, we mapped *petL* transcript 5’ ends by primer extension (Figure 5D). Because *petL* transcripts are considerably less abundant in dPPR-expressing mutants than in the wild-type (Figure 5C), the amount of input RNA for the primer extension assays was adjusted with the intent of increasing signal strength for the dPPR-expressing samples (see details in Figure 5 legend). The wild-type RNA produced a single product of the expected length (109 nucleotides, transcript 1), and this was missing in the *pgr3* mutant as expected. RNA extracted from four different pools of *pgr3* mutants expressing dPPR^*petL*^ contained a novel shorter transcript whose 5’ end mapped ~6 nucleotides downstream of the native end (marked with an asterisk). Because dPPR^*petL*^ was targeted to a sequence internal to the PGR3 binding site (Figure 5A), the position of the novel terminus matches that expected for the product of exonucleolytic degradation back to a dPPR^*petL*^ barrier (~103 nt, see Figure 5A). These results show that an artificial PPR protein can substitute for PGR3’s *petL* RNA stabilization function. That said, the low abundance of dPPR^*petL*^-stabilized RNA indicates that the artificial protein is a less effective RNA stabilizer than is PGR3. Possible explanations are discussed below.

## DISCUSSION

The PPR motif has been proposed as a promising scaffold for the custom-design of RNA binding proteins with desired RNA sequence specificity for the control of RNA metabolism *in vivo* (20). The diversity of functions held by natural PPR proteins *in vivo* predicted many potential applications of PPR-based tools for the manipulation of RNA metabolism. However, first attempts to recode natural PPR proteins to bind new RNA sequences by targeted amino acid mutagenesis faced serious drawbacks due to irregularities found in the amino acid composition of some PPR repeats and the unpredictability of their binding specificity (6,9,15). In addition, most natural PPR proteins appeared to be insoluble when expressed in heterologous systems which limits the ability to engineer them (7). Synthetic consensus PPR repeats (dPPR) were established to overcome these issues (16–18) but their *in vivo* functionalities have just started to be investigated (21).

MRL1 and PGR3 are “pure” (P-type) chloroplastic PPR proteins that harbor tracts of canonical 35 amino acid PPR motifs and lack any accessory domains. MRL1 and PGR3 consist of 10 and 27 PPR repeats, respectively (22,41). Whereas MRL1 controls the stability of processed *rbcL* mRNAs in Arabidopsis, PGR3 plays dual functions in the stabilization of *petL* and *rpl14* mRNAs and stimulation of *petL* and *rps8* translation (23,41). MRL1 and PGR3 bind untranslated mRNA segments and their RNA footprints are represented by abundant sRNA of ~20-30 nucleotides that accumulate *in vivo* (Figure 6). However, like many other P-type PPR proteins, some PPR motifs in MRL1 and PGR3 lack canonical amino acids at the specificity-determining positions which complicate the understanding of their sequence binding specificity according to the PPR code (Figure 6). Similarly to what has been reported for the best-characterized PPR protein, PPR10 (15), some of these motifs might still be critical for the high-affinity interaction of these proteins with their RNA binding site.

**Figure 6.**
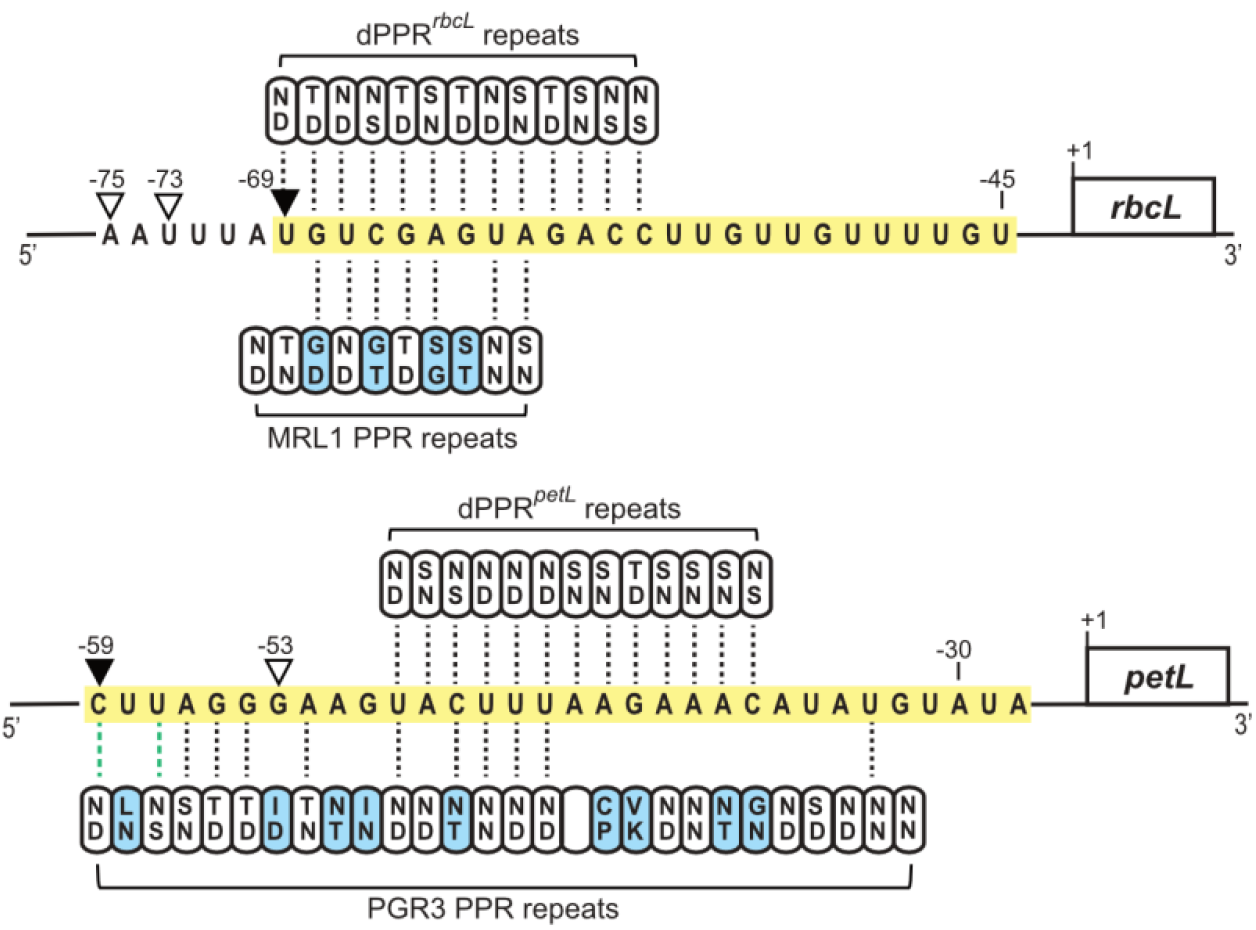
Alignments between dPPR^*rbcL*^, dPPR^*petL*^ and MRL1, PGR3 and their respective designated or predicted *rbcL* and *petL* binding sites. The MRL1 and PGR3 *in vivo* RNA footprints are highlighted in yellow. The MRL1- and PGR3- or dPPR^*rbcL*^- and dPPR^*petL*^-dependent *in vivo* 5’ ends are indicated by black or white arrowheads, respectively. The specificity-determining amino acids in each PPR motif (positions 5 and 35) are indicated. The PPR motifs that exhibit canonical amino acid combinations according to the original PPR code (9) are colored in white while the others are in blue. PPR repeats whose amino acid combinations match the nucleotide binding prediction according to a recently extended PPR code (11) are indicated with dotted black lines. Dotted green lines indicate compatible PPR nucleotide binding but with low affinity.

Our functional complementation assays of Arabidopsis mutants lacking MRL1 and PGR3 by dPPRs built from 13 synthetic PPR repeats add to the evidence that dPPRs bind their designated RNA targets with specificity *in vivo*, and additionally revealed that they can protect RNA downstream from their binding site against the action of exoribonucleases. Similarly to the action of native PPR proteins, this dPPR-dependent RNase blockade mechanism allowed the definition of novel 5’-end processed mRNAs in chloroplasts and control of RNA stability. Therefore, our study demonstrated that artificial dPPRs hold intrinsic properties that are functionally equivalent to natural PPRs. Nevertheless, the dPPR^*rbcL*^ and dPPR^*petL*^ experimental designs revealed interesting differences in their efficiency to stabilize their cognate mRNA targets compare to their natural PPR counterparts. Whereas dPPR^*rbcL*^ fully complemented the *rbcL* mRNA stabilization function of MRL1, dPPR^*petL*^ only partially stabilized its cognate *petL* mRNA ligand as compared to PGR3. The relatively poor RNA stabilization by dPPR^*petL*^ most likely results from the fact that it contains fewer than half of the PPR motifs as PGR3 (13 *versus* ~30 motifs) (Figure 6): this provides many fewer protein-RNA contacts (and thus presumably lower RNA binding affinity) and may also leave endonuclease-sensitive RNA sequences exposed that are masked by PGR3. By contrast, dPPR^*rbcL*^ harbors more PPR repeats than its counterpart MRL1 (Figure 6). Alternatively, this difference may result simply from different expression levels or from inefficient chloroplast targeting due to the use of a maize transit peptide on dPPR^*petL*^.

In conclusion, our results demonstrate that synthetic PPR proteins can be used for the targeted stabilization of specific chloroplastic mRNAs and therefore, the manipulation of organellar gene expression. Finally, this study provides knowledge upon which to develop strategies for the regulation of RNA turn-over in plant organelles by the use of designer PPRs.

## AVAILABILITY

The NGS data discussed in this publication have been deposited in NCBI’s Gene Expression Omnibus (44) and are accessible through GEO Series accession number GSE146249 (https://www.ncbi.nlm.nih.gov/geo/query/acc.cgi?acc=GSE146249).

## ACCESSION NUMBERS

Sequence data from this article can be found in the EMBL/GenBank libraries under the following accession numbers: *MRL1* (At4g34830) and *PGR3* (At4g31850) from A. thaliana, A. thaliana chloroplast genome (NC_000932.1).

## SUPPLEMENTARY DATA

**Supplementary Figure 1.** DNA and protein sequences of dPPR^*rbcL*^ and dPPR^*petL*^

**Supplementary Figure 2.** RIP-seq data replicates

**Supplementary Figure 3.** Closer view of *rbcL* RNA peak in the RIP-seq analysis.

**Supplementary Figure 4.** Phenotype of dPPR^*petL*^ transformants.

**Supplementary Table 1.** List of oligonucleotides

**Supplementary Table 2.** RNA peaks in dPPR^*rbcL*^ RIP-seq analysis Supplementary Data are available at NAR online.

## ACKNOWLEDGEMENT

We are grateful to Olivier Vallon (CNRS UMR7141) for the gift of *mrl1* seeds and Toshiharu Shikanai (Kyoto University) for the gift of *pgr3* seeds and NdhL antibody.

## FUNDING

This work was supported by Agence National de la Recherche [ANR-18-CE20-0013 to K.H.]; the European Union’s Horizon 2020 research and innovation programme under the Marie Sklodowska-Curie action [grant number 794377 to N.M.]; and U.S.-Israel Binational Agricultural Research and Development Grant [IS-5205-19 to A.B.]; Funding for open access charge: Centre National de la Recherche Scientifique.

## CONFLICT OF INTEREST

We have no conflict of interest to declare.

---

**Supplementary Figure 1.** Sequences of dPPR^*rbcL*^ and dPPR^*petL*^. RecA and PPR10 transit peptides are highlighted in green, PPR10-derived sequences in blue and the artificial PPR tracts are shown in black. The dPPR^*rbcL*^ C-terminal 3xHA tag is brought by the binary vector and is not shown in the sequences. The dPPR^*petL*^ C-terminal FLAG tag is shown in bold.

**Supplementary Figure 2. RIP-seq data replicates.** The RIP-seq analysis for the experimental (dPPR^*rbcL*^) and control (Col-0) immunoprecipitations were as described in Figure 4B. The graphs plot the kiloreads per nucleotide along the entire Arabidopsis chloroplast genome (NC_000932.1). Due to high coverage, the results corresponding to the chloroplast rRNAs loci (100,001-154,478) are displayed in an independent graph on the right.

**Supplementary Figure 3.** Closer view of *rbcL* RNA peak in the RIP-seq analysis from Figure 4B.

**Supplementary Figure 4.** Phenotypes of *pgr3-4*:dPPR^*petL*^ plants. Plants were grown for 24 days at 22°C in short day conditions (70 μE light intensity).

**Supplementary Table 1.** List of oligonucleotides.

**Supplementary Table 2.** RNA peaks in dPPR^*rbcL*^ RIP-seq analysis.

## Notes

### Competing Interest Statement

The authors have declared no competing interest.

