## Supplementary for "*In vivo* stabilization of endogenous chloroplast RNAs by customized artificial pentatricopeptide repeat proteins"

### DNA sequence of *dPPR*$^{rbcL}$

ATGGATTCTCAACTTGTTCTTTCTCTTAAGCTTAACCCTTCTTTTACTCCTCTTTCTCCTCTTTTTCCTTTTACTCCTTGTTC
TTCTTTTTCTCCTTCTCTTAGATTTTCTTCTTGTTATTCTAGGAGACTTTACTCTCCTGTTACTGTTTATGCTGCTAAGAAGC
TTTCTCATAAGATTTCTTCTGAGTTTGATGATAGAATCTCTTTGCCTCTTGATTCTTTGCTTCTTCATCTTACTGCTCCTGCT
CCTGCTCCAGCTCCTGCTCCTAGAAGATCTCATCAAACTCCTACTCCTCCTCATTCTTTTCTTTCTCCTGATGCTCAAGTTCT
TGTTCTTGCTATTTCTTCTCATCCTCTTCCTACTCTTGCTGCTTTTCTTGCTTCTAGGAGAGATGAACTTCTTAGAGCTGATA
TTACCTCTCTTCTTAAGGCTTTGGAACTCTCTGGACATTGGGAATGGGCTCTTGCTCTTCTTAGATGGGCTGGAAAGGAGGGA
GCTGCTGATGCTTCTGCTCTTGAAATGGTTGTTAGGGCATTGGGAAGAGAAGGTCAGCATGATGCTGTTTGTGCTTTGTTGGA
TGAAACTCCTCTTCCTCCTGGATCTAGGCTTGATGTTAGAGCTTATACTACTGTTTTGCATGCTTTGAGTAGAGCTGGTAGAT
ATGAAAGGGCTTTGGAATTGTTTGCTGAGCTTAGAAGGCAAGGAGTTGCTCCTACTGTTGTTACTTATAATACTCTTATCGAT
GGACTTTGTAAGGCTGGTAAGTTGGATGAGGCACTTAAGTTGTTTGAAGAAATGGTGGAAAAGGGAATAAAACCAGATGTTGT
TACTTACACTACTCTTATTGATGGACTTTGCAAGGCTGGAAAGTTGGATGAAGCACTTAAGCTTTTTGAAGAAATGGTTGAAA
AGGGAATTAAGCCTGATGTTGTTACATATAATACCCTCATTGATGGATTGTGTAAGGCTGGAAAATTGGATGAAGCTCTTAAG
TTGTTCGAGGAAATGGTGGAGAAGGGAATTAAACCTGATGTTGTGACTTATAATACCCTTATTGATGGTCTTTGTAAGGCAGG
TAAGTTGGATGAAGCTTTGAAGTTGTTTGAGGAAATGGTTGAGAAGGGAATCAAGCCATCTGTTGTTACTTATACTACTCTTA
TCGATGGTCTTTGCAAGGCAGGAAAGTTGGATGAAGCTCTCAAGTTGTTTGAAGAGATGGTTGAAAAAGGAATTAAGCCAGAT
GTTGTGACATATAGTACTCTTATTGATGGACTCTGCAAGGCTGGTAAATTGGATGAGGCTCTTAAGTTATTTGAAGAGATGGT
GGAGAAAGGAATTAAACCAAATGTTGTGACTTACACTACACTTATTGATGGATTATGTAAGGCAGGAAAACTTGATGAAGCTT
TAAAGTTGTTTGAGGAGATGGTGGAAAAAGGAATCAAGCCTGATGTGGTGACATATAATACTTTAATTGATGGACTCTGTAAG
GCTGGAAAGCTCGATGAAGCTCTTAAATTGTTTGAGGAAATGGTTGAAAAGGGTATTAAGCCTGATGTTGTTACCTATTCTAC
TTTGATTGATGGTTTGTGTAAGGCAGGAAAGTTAGATGAAGCATTGAAGTTGTTCGAAGAAATGGTTGAAAAGGGAATCAAAC
CAAATGTGGTGACTTATACTACACTTATCGATGGATTGTGCAAGGCTGGAAAGTTAGATGAGGCTTTGAAGTTATTTGAGGAA
ATGGTTGAAAAGGGAATTAAGCCTGATGTTGTTACTTATTCTACTCTTATTGATGGACTTTGTAAAGCAGGTAAGCTCGATGA
GGCATTGAAGTTATTCGAGGAAATGGTTGAAAAGGGAATTAAGCCTAATGTTGTTACTTATAACACCCTTATTGATGGACTTT
GTAAGGCTGGAAAGCTTGATGAGGCATTAAAGTTGTTCGAAGAGATGGTTGAGAAAGGAATAAAGCCTTCTGTTGTGACTTAT
AACACTTTGATTGATGGACTTTGTAAGGCTGGAAAGCTTGATGAAGCACTCAAGTTGTTCGAAGAAATGGTTGAAAAGGGAAT
TAAGCCTTCAGAATTGACTTATAGGAGGGTTGTTGAAAGTTATTGTAGGGCTAAAAGATTTGAAGAGGCTAGAGGATTTCTTT
CTGAAGTTAGTGAAACTGATTTGGATTTCGATAAGAAGGCTTTGGAGGCTTATATTGAGGATGCTCAGTTTGGTAGG

### Amino acid sequence of dPPR$^{rbcL}$

MDSQLVLSLKLNPSFTPLSPLFPFTPCSSFSPSLRFSSCYSRRLYSPVTVYAAKKLSHKISSEFDDRISLPLDSLLLHLTAPA
PAPAPAPRRSHQTPTPPHSFLSPDAQVLVLAISSHPLPTLAAFLASRRDELLRADITSLLKALELSGHWEWALALLRWAGKEG
AADASALEMVVRALGREGQHDAVCALLDETPLPPGSRLDVRAYTTVLHALSRAGRYERALELFAELRRQGVAPTVVTYNTLID
GLCKAGKLDEALKLFEEMVEKGIKPDVVTYTTLIDGLCKAGKLDEALKLFEEMVEKGIKPDVVTYNTLIDGLCKAGKLDEALK
LFEEMVEKGIKPDVVTYNTLIDGLCKAGKLDEALKLFEEMVEKGIKPSVVTYTTLIDGLCKAGKLDEALKLFEEMVEKGIKPD
VVTYSTLIDGLCKAGKLDEALKLFEEMVEKGIKPNVVTYTTLIDGLCKAGKLDEALKLFEEMVEKGIKPDVVTYNTLIDGLCK
AGKLDEALKLFEEMVEKGIKPDVVTYSTLIDGLCKAGKLDEALKLFEEMVEKGIKPNVVTYTTLIDGLCKAGKLDEALKLFEE
MVEKGIKPDVVTYSTLIDGLCKAGKLDEALKLFEEMVEKGIKPNVVTYNTLIDGLCKAGKLDEALKLFEEMVEKGIKPSVVTY
NTLIDGLCKAGKLDEALKLFEEMVEKGIKPSELTYRRVVESYCRAKRFEEARGFLSEVSETDLDFDKKALEAYIEDAQFGR

### DNA sequence of *dPPR*$^{petL}$

ATGGAGGCTACAGGTCGAGGTCTCTTTCCTAATAAGCCTACGCTTCCTGCAGGACCTAGAAAGCGTGGCCCTCTCCTCCCAGC
TGCTCCGCCGCCTCCAAGTCCCAGTAGTCTGCCATTGGATTCTCTATTGCTGCATTTGACAGCGCCGGCTCCTGCACCTGCAC
CGGCACCAAGACGATCCCATCAGACACCAACCCCACCTCACTCTTTCCTGTCTCCAGACGCTCAGGTGTTGGTCCTTGCAATA
AGCTCTCACCCGCTTCCGACACTTGCGGCATTCTTGGCTTCTAGGCGTGATGAACTATTGAGAGCTGACATTACATCATTGCT
TAAGGCCTTGGAATTGTCTGGCCACTGGGAATGGGCGTTGGCGTTACTCAGATGGGCGGGAAAGGAAGGAGCTGCTGATGCAT
CCGCATTGGAAATGGTGGTCAGAGCTTTGGGAAGGGAGGGTCAACATGATGCTGTGTGTGCTCTCCTTGATGAAACTCCTCTT
CCTCCTGGGTCTCGTCTTGATGTTCGTGCATATACGACAGTTTTACATGCTCTGTCGAGGGCGGGAAGGTACGAGCGAGCCTT
AGAATTGTTCGCCGAATTAAGGCGTCAGGGAGTGGCTCCTACTGTTGTTACGTATAACACCTTGATCGATGGACTGTGTAAGG
CCGGGAAGCTAGATGAAGCTCTTAAATTATTCGAAGAGATGGTGGAGAAGGGTATCAAGCCAGACGTCGTAACATATTCCACC
CTAATCGATGGTCTGTGCAAGGCAGGTAAACTCGATGAAGCCCTAAAATTGTTTGAGGAAATGGTAGAGAAAGGTATTAAACC
GAATGTGGTGACGTACAATACTCTCATCGATGGTCTTTGCAAAGCTGGGAAGTTGGACGAGGCTCTGAAATTGTTTGAAGAGA
TGGTAGAAAAAGGCATAAAACCATCCGTGGTCACTTATAACACACTTATAGACGGCTTGTGTAAAGCTGGGAAGTTGGACGAA
GCTCTAAAGCTATTCGAGGAAATGGTTGAGAAAGGAATTAAGCCAGACGTGGTTACCTACAATACGCTGATCGATGGCCTATG
TAAAGCCGGGAAACTTGATGAAGCACTTAAACTCTTCGAGGAGATGGTGGAAAAAGGAATCAAGCCCGATGTTGTCACATACA
ATACGCTAATTGACGGCCTCTGTAAGGCGGGAAAGCTGGATGAGGCTTTAAAACTTTTCGAAGAGATGGTGGAGAAAGGAATC
AAACCGGACGTTGTAACGTATTCAACTTTAATCGACGGGCTATGCAAGGCTGGGAAGCTCGATGAGGCCTTGAAGTTATTTGA
AGAAATGGTGGAAAAAGGGATCAAACCAAATGTGGTTACTTACTCTACTCTCATCGATGGTCTCTGTAAGGCAGGGAAACTTG
ATGAGGCACTAAAGCTCTTCGAAGAGATGGTGGAGAAAGGTATAAAGCCTAACGTAGTTACGTACACTACGCTGATCGACGGC
TTGGCAAAGTGTGGAAAACTCGACGAGGCACTTAAACTCTTTGAAGAGATGGTCGAAAAGGGGATAAAGCCTGACGTCGTTAC
ATATTCTACGCTAATCGACGGCTTGTGCAAAGCGGGAAAACTAGATGAGGCTTTGAAACTCTTTGAGGAGATGGTTGAGAAGG
GAATCAAGCCGAATGTGGTTACCTACTCAACATTGATCGACGGCCTTTGTAAAGCTGGTAAACTGGATGAAGCACTGAAACTT
TTCGAAGAGATGGTCGAGAAAGGCATAAAGCCAAACGTAGTTACCTATTCAACGCTTATAGATGGGCTGTGCAAAGCCGGTAA
GTTGGATGAAGCCTTAAAACTATTTGAGGAGATGGTAGAGAAGGGTATAAAGCCAAATGTCGTGACGTATAATACGCTAATCG
ATGGGCTTTGCAAGGCAGGTAAACTAGATGAGGCTCTGAAGCTCTTTGAGGAAATGGTTGAAAAGGGAATTAAACCTTCGGAA
CTCACCTACCGACGAGTGGTCGAGAGCTATTGTCGTGCCAAGCGTTTCGAGGAAGCAAGAGGCTTTTTATCTGAGGTCTCAGA
GACAGACCTTGATTTCGATAAGAAGGCACTGGAAGCCTACATAGAGGACGCTCAATTCGGACGTGGTACC**ATGGACTACAAAG
ACCATGATGGAGACTATAAGGATCACGACATCGATTACAAGGACGATGACGATAAGTAA**

**Amino acid sequence of dPPR*petL***

MEATGRGLFPNKPTLPAGPRKRGPLLPAAPPPPSPSSLPLDSLLLHLTAPAPAPAPAPRRSHQTPTPPHSFLSPDAQVLVLAI
SSHPLPTLAAFLASRRDELLRADITSLLKALELSGHWEWALALLRWAGKEGAADASALEMVVRALGREGQHDAVCALLDETPL
PPGSRLDVRAYTTVLHALSRAGRYERALELFAELRRQGVAPTVVTYNTLIDGLCKAGKLDEALKLFEEMVEKGIKPDVVTYST
LIDGLCKAGKLDEALKLFEEMVEKGIKPNVVTYNTLIDGLCKAGKLDEALKLFEEMVEKGIKPSVVTYNTLIDGLCKAGKLDE
ALKLFEEMVEKGIKPDVVTYNTLIDGLCKAGKLDEALKLFEEMVEKGIKPDVVTYNTLIDGLCKAGKLDEALKLFEEMVEKGI
KPDVVTYSTLIDGLCKAGKLDEALKLFEEMVEKGIKPNVVTYSTLIDGLCKAGKLDEALKLFEEMVEKGIKPNVVTYTTLIDG
LAKCGKLDEALKLFEEMVEKGIKPDVVTYSTLIDGLCKAGKLDEALKLFEEMVEKGIKPNVVTYSTLIDGLCKAGKLDEALKL
FEEMVEKGIKPNVVTYSTLIDGLCKAGKLDEALKLFEEMVEKGIKPNVVTYNTLIDGLCKAGKLDEALKLFEEMVEKGIKPSE
LTYRRVVESYCRAKRFEEARGFLSEVSETDLDFDKKALEAYIEDAQFGRGT**MDYKDHDGDYKDHDIDYKDDDDK**

**Supplementary Figure 1.** Sequences of dPPR*rbcL* and dPPR*petL*. RecA and PPR10 transit peptides are highlighted in green, PPR10-derived sequences in blue and the artificial PPR tracts are shown in black. The dPPR*rbcL* C-terminal 3xHA tag is brought by the binary vector and is not shown in the sequences. The dPPR*petL* C-terminal FLAG tag is shown in bold.

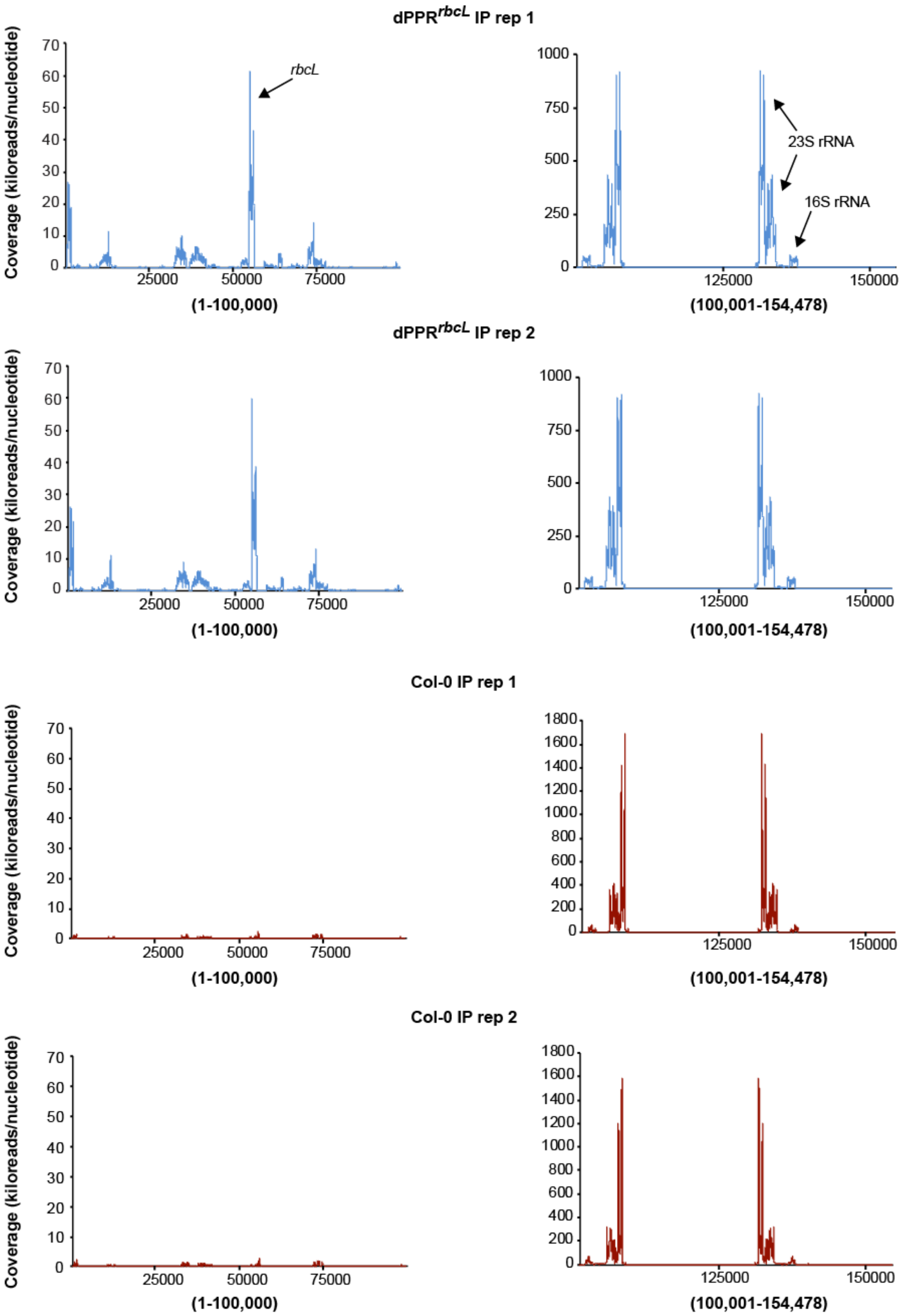


**Supplementary Figure 2. RIP-seq data replicates.** The RIP-seq analysis for the experimental (dPPR$^{rbcL}$) and control (Col-0) immunoprecipitations were as described in Figure 4B. The graphs plot the kiloreads per nucleotide along the entire Arabidopsis chloroplast genome (NC_000932.1). Due to high coverage, the results corresponding to the chloroplast rRNAs loci (100,001-154,478) are displayed in an independent graph on the right.

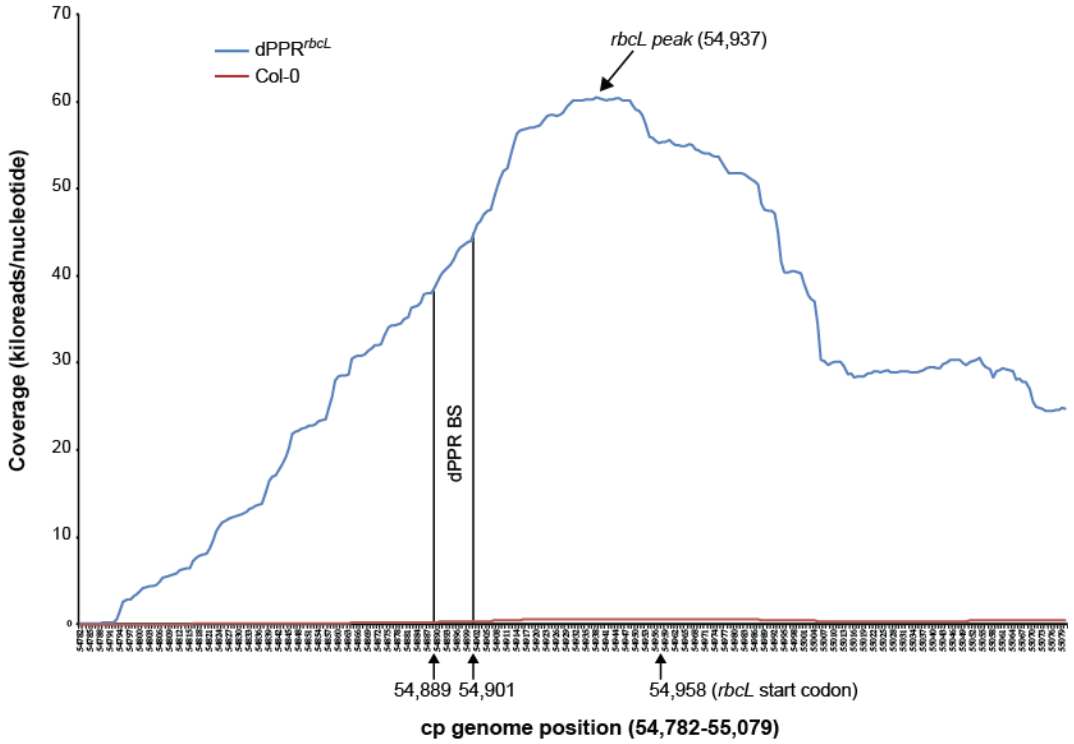


**Supplementary Figure 3.** Closer view of *rbcL* RNA peak in the RIP-seq analysis from Figure 4B.

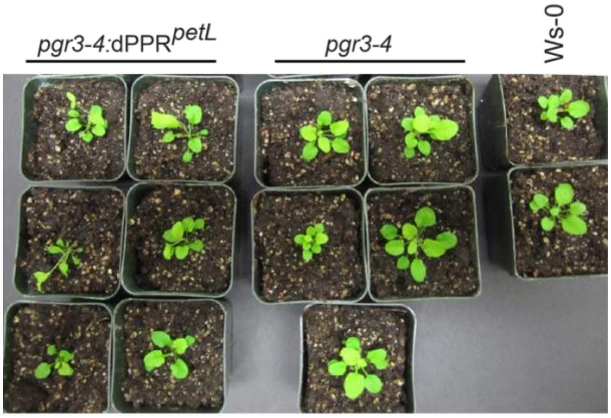


**Supplementary Figure 4. Phenotype of dPPR$^{petL}$ plants.** Plants were grown for 24 days at 22˚C in short day conditions (70 µE light intensity).

**Supplementary Table 1**. List of oligonucleotides

| Name | Sequence 5‘->3‘ | Experiment |
|---|---|---|
| K450Fw | GGGGACAAGTTTGTACAAAAAAGCAGGCTTCATGGATTCTCA ACTTGTTCTTTCTC | dPPR$^{rbcL}$ binary vectors cloning |
| K628Rev | GAAGCAAAGAATCAAGAGGCAAAGAGATTCTATCATCAAACT CAGAAGAAATC | dPPR$^{rbcL}$ binary vectors cloning |
| K627Fw | GATTTCTTCTGAGTTTGATGATAGAATCTCTTTGCCTCTTGATT CTTTGCTTC | dPPR$^{rbcL}$ binary vectors cloning |
| K465Rev | GGGGACCACTTTGTACAAGAAAGCTGGGTCCCTACCAAACTG AGCATCCTCAA | dPPR$^{rbcL}$ binary vectors cloning |
| K461Fw | TAGGATCCTCTTTGCCTCTTGATTCTTTGCTTC | pMAL cloning |
| K462Rev | TATGTCGACTTACCTACCAAACTGAGCATCCTCAA | pMAL cloning |
| rbcL 5' UTR PE | GTCCCTCCCTACAAGTCATGA | Primer extension |
| 5' At rbcL cRT | ACTCGGAATGCTGCCAAGAT | cRT-PCR |
| 3' At rbcL cRT | AGCTTGCAAATGGAGTCCTG | cRT-PCR |
| *rbcL*-1 60-mer | ATACTCTTTAACACCAGCTTTGAACCCAACACTTGCTTTAGTC TCTGTTTGTGGTGACAT | RNA gel blot/Slot blot |
| *psbE* 60-mer | GGGCTCCCGAACACATCGTAAGCTAAACCGGTGCTGACGAAT AACCAGCCCGCAATGAAT | RNA gel blot/Slot blot |
| *psbA* 60-mer | GACGGTTTTCAGTGCTAGTTATCCAGTTACAGAAGCGACCCC ATAGGCTTTCGCTTTCGC | Slot blot |
| *atpF* 60-mer | CATCGTACCAAACATCCCAATATTTGCATTAATAGTACGTAAA TGTAACTCATTACTCAA | Slot blot |
| *psbC* 60-mer | CCCGCTGCCGCTGCCCGAGCCCTTCCCGCGTGCCATAAATG ACCCACGAATAGGAAAAAT | Slot blot |
| *psaB* 60-mer | CCTTGCCAAGCTACATGAAACAAATTTCCGGAAGTCCACAGA AAAATTATTGCTAATTGC | Slot blot |
| *psbT* 60-mer | AAAGTGGATACTAAGAGAAATGTATAAACCAATGCTTCCATAA ATTTGATCGTGGTTTAC | Slot blot |
| *petL* 60-mer | GACCAATAAACAGAACTGAGGTTATAGTTAAAGCTGCTAGTA GAAAACCGAAATAACTAG | RNA gel blot |
| *petL*_PE | ATAGTTAAAGCTGCTAGTAG | Primer extension |

**Supplementary Table 2**. RNA peaks in the dPPR*rbcL* RIP-seq analysis (supporting Figure 4B). The peak coverage (reads/nucleotide), genomic and gene positions are shown.

| Gene | Coverage | Genomic position | Position |
|---|---|---|---|
| ***rbcL*** | **60,508** | **54,937** | **5'UTR** |
| *psbA* | 26,682 | 727 | ORF |
| *psbT* | 14,303 | 73,978 | 5'UTR |
| *atpF* | 11,329 | 12,726 | ORF |
| *psbC* | 10,085 | 34,582 | ORF |
| *psaB* | 6,640 | 39,394 | ORF |
| *psbE* | 6,553 | 64,264 | ORF |
